# Centromere evolution in the fungal genus *Verticillium*

**DOI:** 10.1101/2020.06.29.179234

**Authors:** Michael F Seidl, H Martin Kramer, David E Cook, Gabriel Lorencini Fiorin, Grardy CM van den Berg, Luigi Faino, Bart PHJ Thomma

## Abstract

Centromeres are chromosomal regions that are crucial for chromosome segregation during mitosis and meiosis, and failed centromere formation can contribute to chromosomal anomalies. Despite this conserved function, centromeres differ significantly between and even within species. Thus far, systematic studies into the organization and evolution of fungal centromeres remain scarce. In this study, we identified the centromeres in each of the ten species of the fungal genus *Verticillium* and characterized their organization and evolution. Chromatin immunoprecipitation of the centromere-specific histone CenH3 (ChIP-seq) and chromatin conformation capture (Hi-C) followed by high-throughput sequencing identified eight conserved, large (∼150 kb), AT-, and repeat-rich regional centromeres that are embedded in heterochromatin in the plant pathogen *V. dahliae*. Using Hi-C, we similarly identified repeat-rich centromeres in the other *Verticillium* species. Strikingly, a single repetitive element is strongly associated with centromeric regions in some but not all *Verticillium* species. Extensive chromosomal rearrangements occurred during *Verticillium* evolution, yet only a minority could be linked to centromeres, suggesting that centromeres played a minor role in chromosomal evolution. Nevertheless, the size and organization of centromeres differ considerably between species, and centromere size was found to correlate with the genome-wide repeat content. Overall, our study highlights the contribution of repetitive elements to the diversity and rapid evolution of centromeres within the fungal genus *Verticillium*.

**IMPORTANCE:** The genus *Verticillium* contains ten species of plant-associated fungi, some of which are notorious pathogens. *Verticillium* species evolved by frequent chromosomal rearrangements that contribute to genome plasticity. Centromeres are instrumental for separation of chromosomes during mitosis and meiosis, and failed centromere functionality can lead to chromosomal anomalies. Here, we used a combination of experimental techniques to identify and characterize centromeres in each of the *Verticillium* species. Intriguingly, we could strongly associate a single repetitive element to the centromeres of some of the *Verticillium* species. The presence of this element in the centromeres coincides with increased centromere sizes and genome-wide repeat expansions. Collectively, our findings signify a role of repetitive elements in the function, organization and rapid evolution of centromeres in a set of closely related fungal species.

## INTRODUCTION

Centromeres are crucial for reliable chromosome segregation during mitosis and meiosis. During this process, centromeres direct the assembly of the kinetochore, a multi-protein complex that facilitates attachment of spindle microtubules to chromatids (1–3). Failure in formation or maintenance of centromeres can lead to aneuploidy, i.e. changes in the number of chromosomes within a nucleus, and to chromosomal rearrangements (3–5). While these processes have been often associated with disease development (6), they can also provide genetic diversity that is beneficial for adaptation to novel or changing environments (7, 8). For example, aneuploidy in the budding yeast *Saccharomyces cerevisiae* can lead to increased fitness under selective conditions, such as the presence of antifungal drugs (9, 10). Thus, centromeric instability can contribute to adaptive genome evolution (11, 12).

Despite their conserved function, centromeres are among the most rapidly evolving genomic regions (13, 14) that are typically defined by their unusual (AT-rich) sequence composition, low gene and high repeat density, and heterochromatic nature (13, 15). Nevertheless, centromeres differ significantly in size, composition, and organization between species (13, 16). Centromeres in *S. cerevisiae* are only ∼125 nucleotides long and are bound by a single nucleosome containing the centromere-specific histone 3 variant CenH3 (also called CENP-A or Cse4) (17–20). In contrast to these ‘point centromeres’, centromeres in many other fungi are more variable and larger, and have thus been referred to as ‘regional centromeres’ (15). For instance, in the opportunistically pathogenic yeast *Candida albicans*, the CenH3-bound 3-5 kb long centromeric DNA regions differ significantly between chromosomes, and rapidly diverged from closely related *Candida* species (21–23). Centromeres in the basidiomycete yeasts *Malassezia* are similar in size (3-5 kb) but contain a short AT-rich consensus sequence in multiple *Malassezia* species (11). In *Malassezia*, chromosomal rearrangements and karyotype changes are driven by centromeric loss through chromosomal breakage or by inactivation through sequence diversification (11). Chromosomal rearrangements at centromeres have been similarly observed in the yeast *Candida parapsilosis*, suggesting that centromeres can be fragile and contribute to karyotype evolution (11, 12). CenH3-bound centromeric regions of the basidiomycete yeast *Cryptococcus neoformans* are relatively large, ranging from 30 to 65 kb, and are rich in Long Terminal Repeat (LTR)-type retrotransposons (16). Centromere sizes differ between *Cryptococcus* species as those lacking RNAi and DNA methylation have shorter centromeres, associated with the loss of full-length LTR retrotransposons at centromeric regions, suggesting that functional RNAi together with DNA methylation is required for centromere stability (16).

In filamentous fungi, centromeres have been most extensively studied in the saprophyte *Neurospora crassa* (15). In this species, centromeric regions are considerably larger than in yeasts (on average ∼200 kb), and are characterized by AT-rich sequences that are degenerated remnants of transposable elements and sequence repeats that lack an overall consensus sequence (15, 24, 25). The increased AT-content and the degenerated nature of transposable elements in the genome of *N. crassa* are the result of a process called repeat-induced <u>p</u>oint mutation (RIP) (15, 26). RIP has been linked to the sexual cycle of ascomycetes and targets repetitive sequences by inducing C to T mutations, preferably at CpA di-nucleotides (26). The AT-rich centromeric regions are bound by CenH3 and enriched in the heterochromatin-specific histone modification histone 3 trimethylation of lysine 9 (H3K9me3) (25). Additionally, H3K9me3 and cytosine methylation occurs at the periphery of the centromeres (25). Alterations in H3K9me3 localization compromise centromeric localization, suggesting that the formation and location of heterochromatin, rather than the DNA sequence itself, is essential for function and localization of centromeres in *N. crassa* (15, 25). However, heterochromatin is not a hallmark for centromeres in all filamentous fungi. Centromeres in the fungal wheat pathogen *Zymoseptoria tritici* are shorter (∼10 kb) and AT-poor, and their presence does not correlate with transposable elements nor with heterochromatin-specific histone modifications such as H3K9me3 or histone 3 trimethylation of lysine 27 (H3K27me3) (27). Thus, even though centromeric function is highly conserved, fungal centromeres differ considerably in size, sequence composition, and organization.

Knowledge on centromeres has been impaired by their repetitive nature, which hampers their assembly and subsequent analyses (15, 28). However, recent advances in long-read sequencing technologies enables to study the constitution and evolution of centromeres (11, 16, 29–31). By using long-read sequencing technologies in combination with optical mapping, we previously generated gapless genome assemblies of two strains of the fungal plant pathogen *Verticillium dahliae* (32), whose genomes are characterized by genome rearrangements and the occurrence of lineage-specific (LS) regions (7, 8, 33–35) that are hypervariable between *V. dahliae* strains and contain genes with roles in adaptive evolution to plant hosts (7, 8, 33, 35). Repetitive elements within the LS regions display a distinct chromatin state when compared with other repetitive regions (36). The *Verticillium* genus consists of ten species that are all soil-borne and presumed asexual but have different life-styles (37). Nine of these species are haploid, while the species *Verticillium longisporum* is an allodiploid hybrid between a strain that is closely related to *V. dahliae* and an unknown *Verticillium* species (37–39). During the evolution of the different *Verticillium* species frequent chromosomal rearrangements occurred (8, 35, 40). Facilitated by the availability of high-quality genome assemblies of *V. dahliae* strains and of all other *Verticillium* species (32, 33, 40, 41), we here sought to identify and study the constitution and evolution of centromeres in the *Verticillium* genus.

## RESULTS

### CenH3-binding identifies large regional centromeres in *Verticillium dahliae*

Centromeres differ significantly between fungi, but most centromeres are functionally defined by nucleosomes containing CenH3 (1). To identify centromeres in *V. dahliae* strain JR2 by chromatin immunoprecipitation followed by high-throughput sequencing (ChIP-seq), we first identified the *V. dahliae* CenH3 ortholog (**Fig. S1a**) and generated transformants with N-terminally FLAG-tagged CenH3 (Table S1). To this end, the coding sequence for the FLAG-tagged CenH3 was inserted in locus behind the native *CenH3* promotor (**Figs. S1b-c**). We subsequently used anti-FLAG antibodies to purify FLAG-tagged CenH3-containing nucleosomes from two *V. dahliae* transformants (**Table S1a**) and sequenced the nucleosome-associated genomic DNA. Mapping of the sequencing reads to the *V. dahliae* strain JR2 genome assembly identified a single CenH3-enriched region per chromosome (**Fig. 1a; Fig. S1d-e**), while mapping of the sequencing reads derived from the WT strain did not reveal any CenH3-enriched region (**Fig. S1d-e**). The CenH3-enriched regions, designated as *Cen1-8*, range between ∼94 and ∼187 kb in size (**Fig. 1a; Table 1**). To corroborate these centromere sizes, we assessed centromere locations based on a previously generated optical map (32, 35) revealing no significant size differences (**Fig. S1f**). Thus, we conclude that CenH3-binding defines large regional centromeres in *V*. *dahliae* strain JR2.

**Figure 1.**
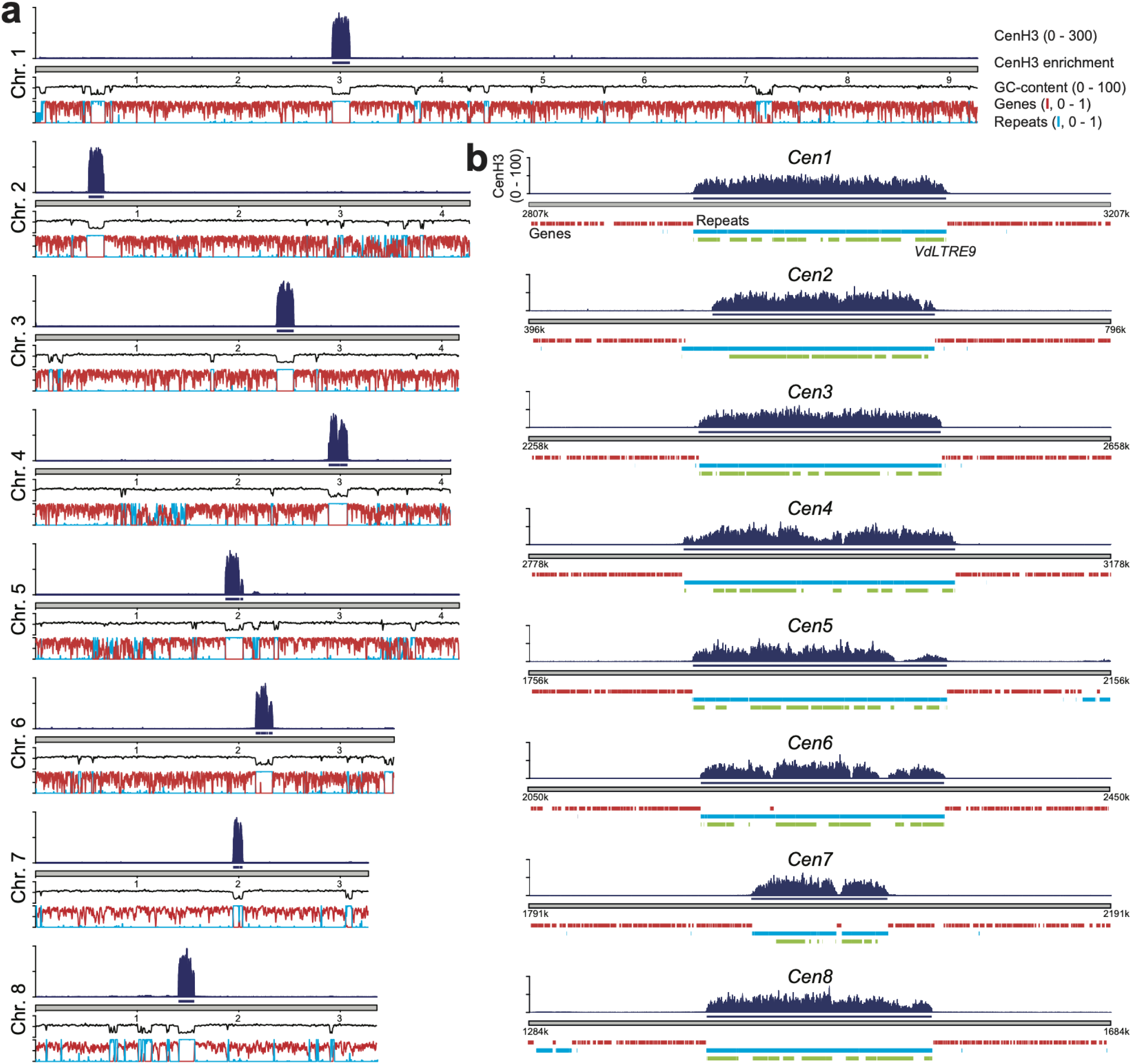
CenH3-binding defines centromeres in *Verticillium dahliae* strain JR2. (a) Schematic overview of the chromosomes of *V. dahliae* strain JR2 showing the normalized CenH3 ChIP-seq read coverage (RPGC normalization in 1 kb bins with 3 kb smoothening), CenH3 enriched regions, GC-content, gene density (red line), and repeat density (blue line). (b) Magnification of a 400 kb region containing the centromere is shown for each of the eight chromosomes of *V. dahliae* strain JR2 (*Cen1-8*) depicting the CenH3 ChIP-seq read coverage (RPGC normalization in 10 bp bins with a 30 bp smoothening) and enrichment, as well as the presence of genes (red) and repetitive elements (blue). Regions carrying the centromere-specific long-terminal repeat element *VdLTRE9* are highlighted in green.

**Table 1:** Genome characteristics of the centromeres of *Verticillium dahliae* strain JR2.

| Chr. | Locus | CenH3 |  | AT-rich | AT-content (%)<sup>3</sup> |  | Repetitive elements |  |
| --- | --- | --- | --- | --- | --- | --- | --- | --- |
|  |  | CenH3 position (bp)<sup>1</sup> | CenH3 length (bp) | Position (kb)<sup>2</sup> | Chr. | Cen. | # Repeats (%)<sup>4</sup> | # VdLTRE9 (%)<sup>4</sup> |
| 1 | *CEN1* | 2,920,143-3,094,179 | 174,037 | 2,919-3,094 | 45.7 | 77.1 | 50 (99.8) | 27 (70.4) |
| 2 | *CEN2* | 520,698-672,281 | 151,584 | 516-672 | 46.3 | 77.8 | 43 (99.7) | 26 (83.0) |
| 3 | *CEN3* | 2,374,294-2,541,026 | 166,733 | 2,375-2,542 | 45.8 | 77.3 | 47 (99.8) | 31 (80.5) |
| 4 | *CEN4* | 2,884,316-3,071,412 | 187,097 | 2,885-3,072 | 46.2 | 75.4 | 54 (99.5) | 24 (53.8) |
| 5 | *CEN5* | 1,868,317-2,043,260 | 174,944 | 1,868-2,044 | 46.7 | 73.9 | 58 (99.5) | 25 (63.1) |
| 6 | *CEN6* | 2,166,972-2,333,060 | 166,089 | 2,167-2,334 | 46.4 | 75.2 | 48 (100) | 31 (62.6) |
| 7 | *CEN7* | 1,944,367-2,038,091 | 93,725 | 1,945-2,038 | 44.7 | 76.5 | 32 (95.8) | 14 (47.8) |
| 8 | *CEN8* | 1,406,398-1,561,664 | 155,267 | 1,406-1,562 | 47.7 | 77.0 | 37 (100) | 26 (73.9) |

<sup>1</sup>: position of CenH3-enriched domains; enriched domains within 10 kb have been merged
<sup>2</sup>: position of AT-rich domains; AT-rich domains with 20 kb have been merged
<sup>3</sup>: average AT-content of 1 kb windows of the entire chromosome and the AT-rich domain
<sup>4</sup>: percentage of centromeric region covered

### Centromeres in *Verticillium dahliae* are repeat-rich and embedded in heterochromatin

Centromeres are often characterized by increased AT-content, increased repeat density, and depletion of protein coding genes (13, 15, 29). To characterize the centromeres in *V. dahliae* strain JR2, we queried the eight chromosomes for the presence of large AT-rich, gene-sparse, and repeat-rich regions. Seven of the eight chromosomes contain only a single large (>93 kb; average size ∼150 kb) AT-rich region (∼74-78% versus ∼46% genome-wide), nearly completely devoid of protein-coding genes and enriched for repetitive sequences, that overlaps with the regions defined by CenH3-binding (**Fig. 1a; Table 1**). In contrast, chromosome 1 contains three regions with these characteristics (**Fig. 1a; Table 1**). However, only one of these overlaps with the centromeric regions defined by CenH3-binding (**Fig. 1**).

Elevated AT-levels in repeat-rich regions are caused by RIP mutations in some filamentous fungi (15, 25, 26, 42). Due to its presumably asexual nature (7), the occurrence of RIP in *V. dahliae* is controversial (8, 43, 44), although signatures of RIP have previously been reported in a subset of repeat-rich regions (36). We assessed the occurrence of RIP signatures in centromeres using the composite RIP index (CRI) (45), which considers C to T mutations in the CpA context. Intriguingly, genomic regions located at centromeres display significantly higher CRI values than other genomic regions (e.g. genes or repetitive elements) (**Fig. 2a; Figs. S2, S3a**), and thus RIP signatures at repetitive elements located at centromeres likely contribute to the high AT-levels.

**Figure 2.**
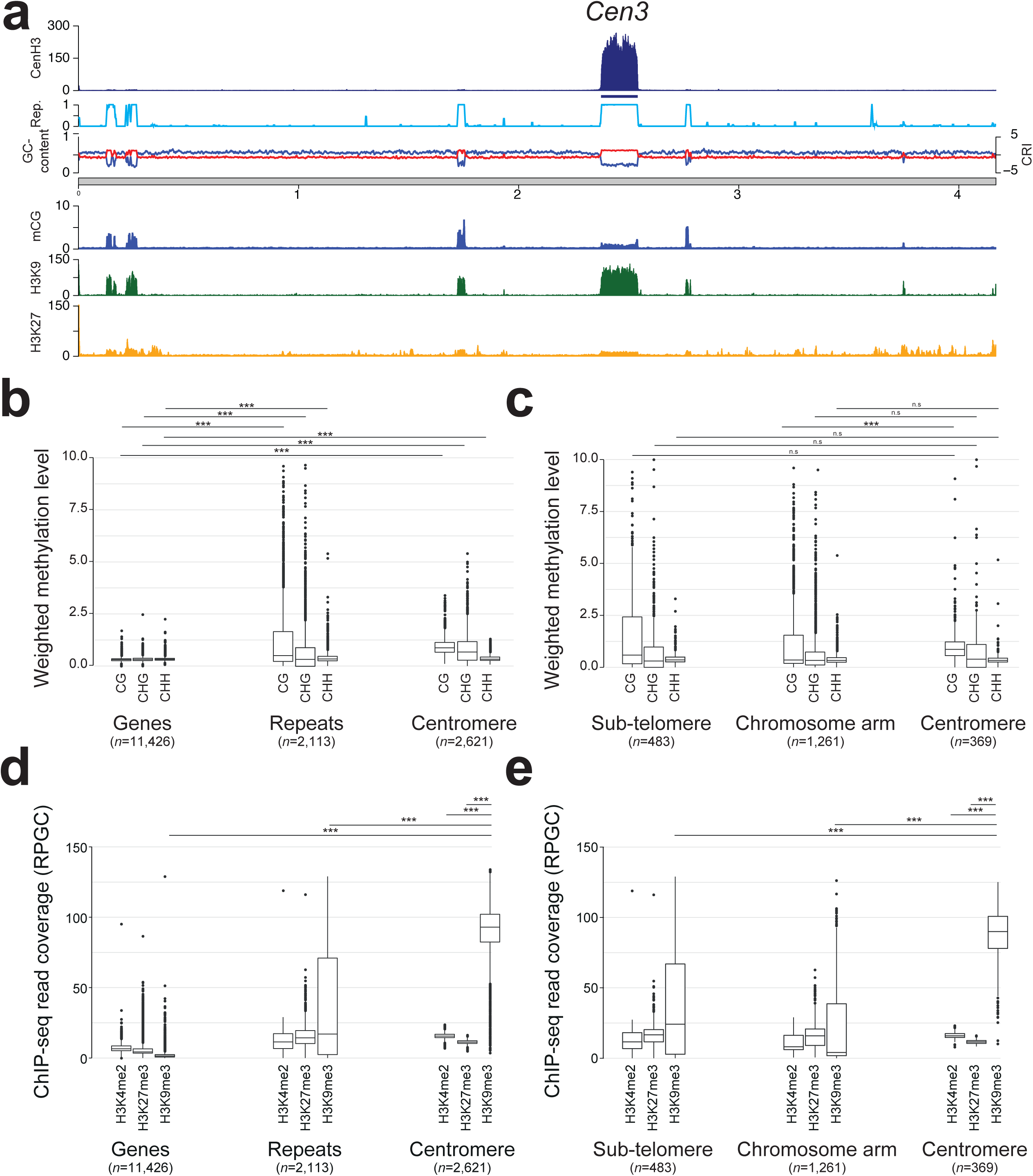
Centromeres in *Verticillium dahliae* strain JR2 are embedded in heterochromatin. (a) Schematic overview of chromosome 3 of *V. dahliae* strain JR2, exemplifying the distribution of heterochromatin-associated chromatin modifications (mC, H3K9me3, and H3K27me3) in relation to the centromeres. The different lanes display the CenH3-FLAG ChIP-seq read coverage (RPGC normalization in 1 kb bins with 3 kb smoothening), the CenH3-FLAG enriched regions, the repeat-density, the GC-content, the CRI as well as the weighted cytosine methylation (all summarized in 5 kb windows with 500 bp slide), and the normalized H3K9me3 and H3K27me3 ChIP-seq read coverage (RPGC normalization in 1 kb bins with 3 kb smoothening). The schematic overview of all chromosomes is shown in Figure S2. (b) Box plots of weighted DNA methylation levels per genomic context (CG, CHG, or CHH) are summarized over genes, repetitive elements, or 5 kb genomic windows (500 bp slide) overlapping with the centromeric regions. (c) Weighted DNA methylation levels per genomic context (CG, CHG, or CHH) are summarized over repetitive elements that have been split based on their genomic location; sub-telomeres (within the first or last 10% of the chromosome), centromeres, or the remainder of the chromosome arm. (d) ChIP-seq read coverage (RPGC normalized; see (a)) for H3K4me2, H3K27m3, and H3K9me3 is summarized over genes, repetitive elements, or 5 kb windows (500 bp slide) overlapping with the centromeric regions. (e) ChIP-seq read coverage (RPGC normalized; see (a)) for H3K4me2, H3K27m3, and H3K9me3 is summarized over repetitive elements that have been split based on their genomic location; sub-telomeres (within the first or last 10% of the chromosome), centromeres, or the remainder of the chromosomal arm. Statistical differences for the indicated comparisons were calculated using the one-sided non-parametric Mann-Whitney test; p-values < 0.001: ***.

In most filamentous fungi and oomycetes, AT- and repeat-rich centromeres are embedded in heterochromatin that is characterized by methylated DNA and by particular histone modifications (H3K9me3 and H3K27me3) (13, 15, 16, 25, 30, 45). We recently determined chromatin states in the genome of *V. dahliae* strain JR2 and revealed that repetitive sequences outside of the LS regions display characteristics of heterochromatin (36). To define centromeric chromatin states, we used previously generated bisulfite sequencing data to monitor DNA methylation (mC) and ChIP-seq data to determine the distribution of the heterochromatic marks H3K9me3 and H3K27me3 (36). To also determine the distribution of euchromatin, we performed ChIP-seq with an antibody against the euchromatic mark di-methylation of lysine 4 of histone H3 (H3K4me2). We observed overall low genome-wide DNA methylation levels (36) (**Fig. 2a; Fig. S2**), similar to the previously reported levels for *Aspergillus flavus* (46) and lower than for *N. crassa* (47). Nevertheless, repetitive elements and centromeres show significantly higher DNA methylation levels in all contexts when compared with genes (**Fig. 2b**). Methylation (in CG context) at repetitive elements at centromeres is significantly higher than at repeats located along the chromosomal arm, but not at sub-telomeric regions (**Fig. 2c**), and more methylation at centromeres correlates with increased CRI (**Fig. 2a; Figs. S2, S3a**). DNA methylation co-localizes with H3K9me3 at repeat-rich regions (36) (**Figs. 2a; Fig. S2**). H3K9me3 occurs predominantly at repetitive elements localized at sub-telomeres and centromeres (**Figs. 2d-e; Figs. S2, S3b**). In comparison, H3K4me2 and H3K27me3 are largely absent from centromeres (**Figs. 2d-e; Fig. S3b**). Collectively, these observations indicate that centromeres of *V. dahliae* display typical characteristics of constitutive heterochromatin.

### A single repeat associates with *Verticillium dahliae* strain JR2 centromeres

Centromere identity and function is typically defined by CenH3-binding and not by specific DNA sequences, although various types of repetitive sequences, such as transposable elements, are commonly observed in centromeres of plants, animals, and fungi (13, 15, 48, 49). Unsurprisingly, CenH3-bound centromeres are repeat-rich in *V. dahliae* (**Fig. 1**). A detailed analysis of the eight centromeres revealed a near-complete (>96%) composition of repetitive elements belonging to only ten different repeat sub-families (**Figs. 1b**, **3a; Table 1**), of which the majority shows similarity to LTR retrotransposons of the *Gypsy*- and *Copia*-like families (**Fig. 3a**). These elements show signs of RIP, are highly methylated, and non-transcribed (**Figs. S3c-e**), and thus likely inactive. Interestingly, a single LTR retrotransposon sub-family, previously designated *VdLTRE9* (8, 32), covers on average ∼70% of the DNA sequences at the eight centromeres, ranging from 47% in *Cen7* to 83% in *Cen2* (**Fig. 3a; Table 1**). We scanned the genome for the localization of the ten repeat sub-families (**Fig. 3**). Intriguingly, although it is one of the most abundant repeats in the genome with 215 complete or partial matches, *VdLTR9* is associated to centromeres as 95% of the copies (204 out of 215; one-sided Fisher’s exact test; multiple-testing corrected p-value 3e-106) occur at the eight centromeres, whereas only 5% of the copies are dispersed over the genome (**Fig. 3b-c**). The nine other repeat sub-families have additional matches that are located outside of the centromeres (**Figs. 1a; Figs. 3b-c**), and only two of these repeats are significantly enriched and consistently present in all eight centromeres; 63% and 45% of the matches of these two sub-families occur at the centromeres (**Fig. 3c**). Collectively, these findings suggest that only the presence of *VdLTRE9* is strongly associated with centromeres in *V. dahliae* strain JR2.

**Figure 3.**
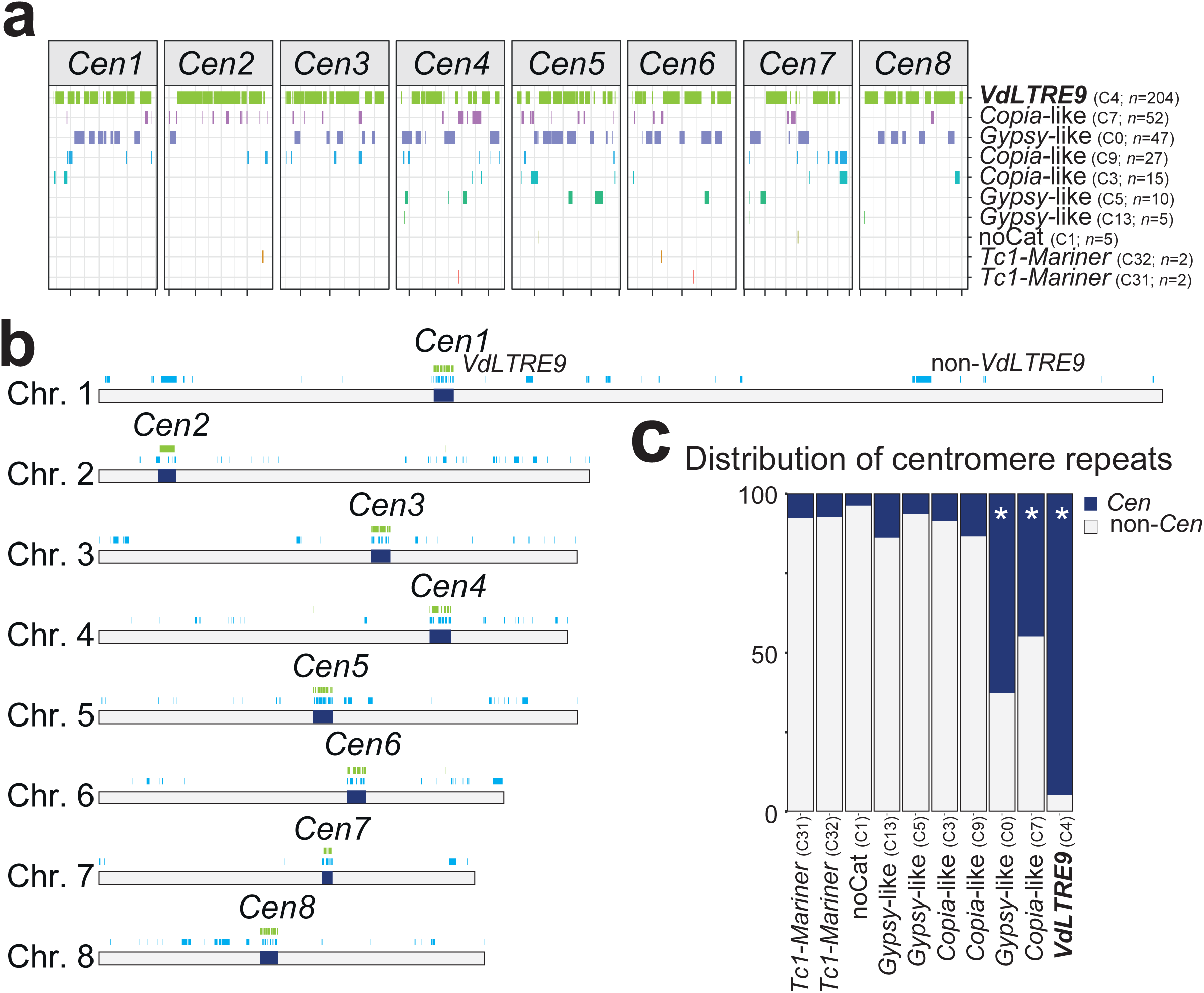
A single repeat family associates with centromeres in *Verticillium dahliae* strain JR2. (a) The presence of different repeat sub-families is shown across the eight centromeres (*Cen1*-*8*), and the number of occurrences for each sub-family within the centromeres is indicated. The individual centromeres in the diagram are shown in equal scale. (b) Genome-wide distribution of the ten repeat sub-families occurring within the eight centromeres (*Cen1*-*8*; dark blue); the location of *VdLTRE9* is shown in green and the location of elements belonging to the other nine sub-repeat families (from panel (a)) is shown in light blue. (c) The distribution of different repeat sub-families in centromeres (*Cen*; dark blue) and across the genome (non-*Cen*; light grey). The enrichment of specific sub-families at centromeres was assessed using a one-sided Fisher’s exact test. Significant enrichment (multiple-testing corrected p-value < 0.01) is denoted with an asterisk.

*VdLTRE9* displays similarity to LTR retrotransposons. The consensus sequence of *VdLTRE9* is ∼7.3 kb long (the two LTR sequences are each ∼200 bp long), and the individual matches share a high degree of sequence identity (∼86%). Sequence similarity based TE-classifications using PASTEC (50) indicates that the consensus sequence displays remote similarity to *Gypsy*-like retrotransposons. Only ∼25% of the *VdLTRE9* matches in the genome cover the entire (>97.5%) consensus sequence, but many of these are still fragmented as they occur as discontinuous copies. Furthermore, the *VdLTRE9* consensus sequence is AT-rich (∼75% AT), which may be caused by RIP (**Fig. S3d**), indicating that *VdLTRE9* has significantly degenerated.

### *VdLTRE9* as hallmark of *Verticillium dahliae* centromeres

To examine if *VdLTRE9* similarly occurs at centromeres in other *V. dahliae* strains, we made use of the complete genome assembly of *V. dahliae* strain VdLs17 (8, 32, 35). The evolution of *V. dahliae* is characterised by chromosomal rearrangements (8, 35) (**Figs. 4a; Figs. S4a-c**). Nevertheless, synteny analyses between *V. dahliae* strains JR2 and VdLs17 revealed large regions of co-linearity between chromosomes and identified significant sequence and synteny conservation between the centromeres and their flanking regions (**Figs. 4b-c; Fig. S4a**), suggesting that centromeric sequences and their locations are conserved. We queried the genome of *V. dahliae* strain VdLs17 for the presence of *VdLTRE9* and identified a single region on each chromosome, collectively containing 186 of the 207 (90%) complete or partial matches of *VdLTRE9* in the genome (**Fig. 4d**) (one-sided Fisher’s exact test; multiple-testing corrected p-value 3e-146). These *VdLTR9*-rich regions are ∼150 kb in size, AT-rich, gene-poor and repeat-rich, and share similarity to the previously identified CenH3-bound and *VdLTRE9*-enriched regions of *V. dahliae* strain JR2 (**Figs. 4b-c; Fig. S4d**), suggesting that these regions similarly represent the centromeres of *V. dahliae* strain VdLs17.

**Figure 4.**
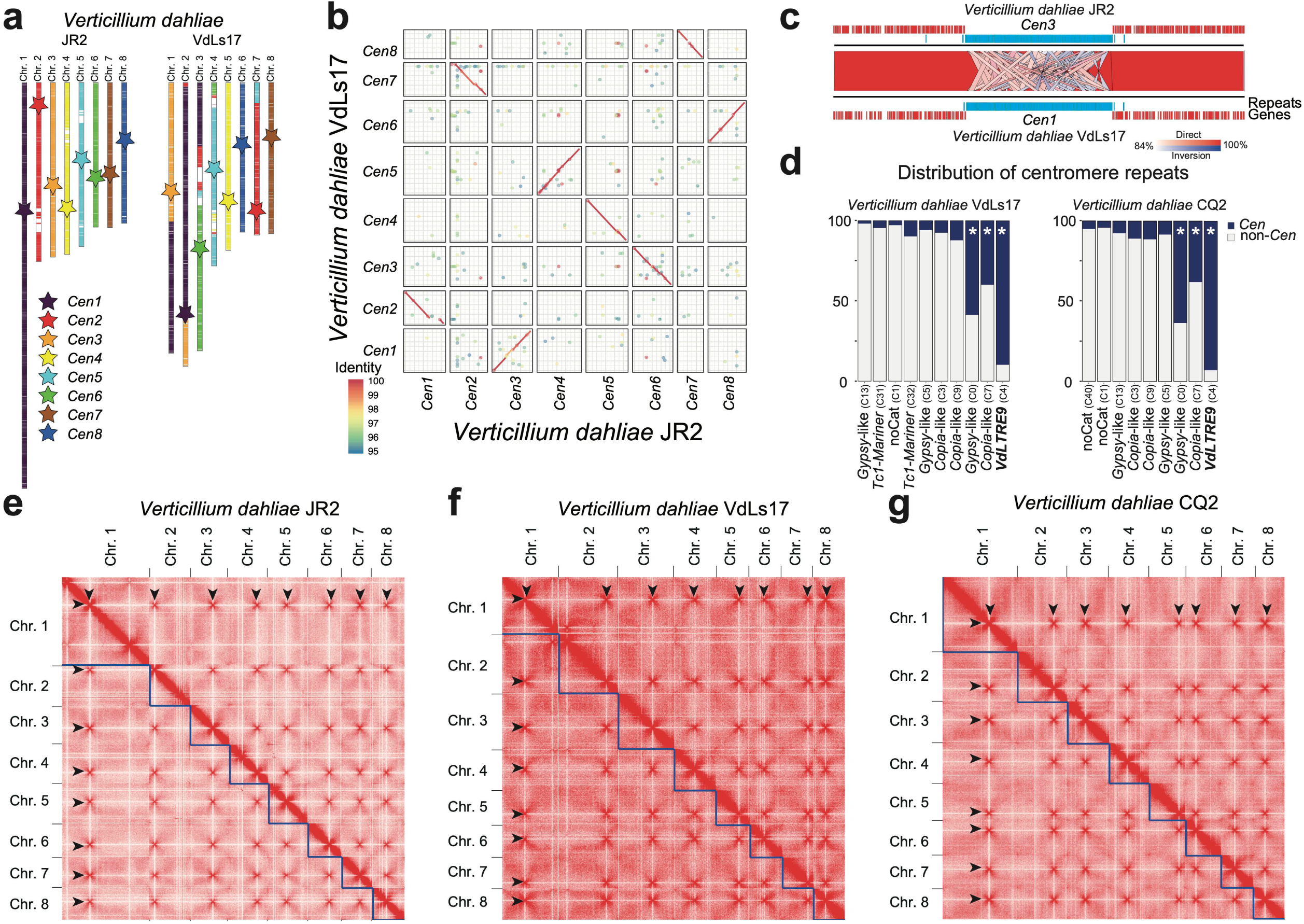
Hi-C contact maps identify *VdLTRE9* as hallmark of centromeres in *Verticillium dahliae*. (a) Synteny analyses of the eight chromosomes of *V. dahliae* strains JR2 and VdLs17. Schematic overview of the eight chromosomes of *V. dahliae* strain JR2 (left) and the corresponding syntenic regions in *V. dahliae* strains VdLs17 (right). Approximate locations of centromeres are indicated by stars, and syntenic centromeres of *V. dahliae* strain VdLs17 are colored according to *Cen1*-*8* of *V. dahliae* strain JR2. (b) Sequence alignment of the centromeric regions ± 20 kb in *V. dahliae* strain JR2 and the corresponding regions in *V. dahliae* strains VdLs17 shown as dot-plot. For clarity, only alignments with >95% sequence identity are displayed. (c) Magnification of *Cen3* of *V. dahliae* strain JR2 and the syntenic *Cen1* of strain VdLs17. Synteny between regions is indicated by ribbons; entire centromeric regions *Cen1* and *Cen3* are syntenic and sequence similarity between individual *VdLTRE9* elements is visualized. The *Cen* regions ± 150 kb are shown as well as genes (red) and repeats (blue) are annotated within this region. (d) Distribution of different repeat families in centromeres (*Cen*; dark blue) and across the genome (non-*Cen*; light grey) for *V. dahliae* strains VdLs17 and CQ2. The enrichment of specific sub-families at centromeres was assessed using a one-sided Fisher’s exact test. Significant enrichment (multiple-testing corrected p-value < 0.01) is denoted with an asterisk. (e-g) Hi-C contact matrix showing interaction frequencies between genomic regions in *Verticillium dahliae* strains JR2 (e), VdLs17 (f), and CQ2 (g). Regions of high inter-chromosomal interaction frequencies are indicative of centromeres and are highlighted by arrow heads. Interaction frequencies are summarized in 50 kb bins along the genome.

Centromeres *N. crassa* and some other fungi co-localize within the nucleus (15, 51–55). This co-localization can be experimentally determined using chromosome conformation capture (Hi-C), which can identify centromeres by their increased inter-chromosomal contacts (55). To confirm that Hi-C can be used to identify centromeres in *V. dahliae*, we first applied Hi-C to *V. dahliae* strain JR2. As anticipated, we observed seven strong inter-chromosomal contacts for each of the eight chromosomes (**Figs. 4e**). Importantly, the interacting regions overlap with the CenH3-bound regions that we identified as centromeres (**Table S1b**), demonstrating that centromeres in *V. dahliae* strain JR2 co-localize within the nucleus and supporting that Hi-C reliably identifies centromeres (51, 52). We then applied Hi-C to *V. dahliae* strain VdLs17, and similarly identified regions with strong inter-chromosomal contacts, one for each of the chromosomes (**Figs. 4f**). These regions overlap with the *VdLTRE9*-enriched regions (**Table S1b**), suggesting that these represent functional centromeres in *V. dahliae* strain VdLs17.

The two *V. dahliae* strains JR2 and VdLs17 are closely related and differ only by ∼0.05% sequence diversity (8, 35). Thus, the conservation of *VdLTRE9* at centromeres could be driven by limited divergence between the two *V. dahliae* strains rather than representing a hallmark of *V. dahliae* centromeres. Therefore, we sought to determine centromeres in an additional *V. dahliae* strain with increased sequence diversity when compared with *V. dahliae* strains JR2 or VdLs17, namely strain CQ2 that displays ∼1.05 percent sequence diversity (33). We previously obtained a long-read based genome assembly of this strain that encompasses 17 contigs (33). We generated Hi-C data for *V. dahliae* strain CQ2 and utilized intra-chromosomal contacts to assign the contigs into eight pseudo-chromosomes, leaving ∼148 kb unplaced scaffolds (**Fig. 4g; Fig. S4e; Table S1c**). We subsequently identified a single region with seven strong inter-chromosomal contacts for each pseudo-chromosome that is significantly enriched for *VdLTRE9* (one-sided Fisher’s exact test; multiple-testing corrected p-value 3.4e-166) (**Figs. 4d, g; Fig. S4e; Table S1b**). Synteny analyses between *V. dahliae* strains JR2 and CQ2 revealed that the eight *VdLTRE9*-rich regions and their flanking chromosomal regions are co-linear, suggesting that centromere locations are conserved between different *V. dahliae* strains (**Figs. 4; Figs. S4a-c, f**). With an average size of 165 kb, the centromeres of *V. dahliae* strain CQ2 are similar in size as the 144 kb and 157 kb average sizes in *V. dahliae* strains VdLs17 and JR2, respectively (**Table S1b**). The sizes of the corresponding (i.e. homologous) centromeres vary between the different *V. dahliae* strains. Yet, the consistent co-occurrence of the *VdLTRE9*-rich regions with the interaction data obtained by Hi-C throughout a selection of *V. dahliae* strains demonstrates that *VdLTRE9* is a hallmark of *V. dahliae* centromeres.

### The evolution of *Verticillium* centromeres

In addition to *V. dahliae*, we previously generated genome assemblies of the eight haploid *Verticillium* species and the allodiploid *V. longisporum* (39, 40) (**Fig. 5a**) that ranged from 12 to 684 scaffolds (**Table S1c**). These ten *Verticillium* species have been traditionally separated over two distinct clades; Flavnonexudans and Flavexudans (**Fig. 5a**) (37). We generated Hi-C data to study the composition and evolution of centromeres in the different *Verticillium* species. By using intra-chromosomal interaction signals, we first assigned the vast majority of the previously assembled contigs into eight pseudo-chromosomes for each of the haploid *Verticillium* species and 16 pseudo-chromosomes for the diploid *V. longisporum*, leaving between 0.5 kb and 2,022 kb unassigned (**Fig. S5; Table S1c**). For most genome assemblies, the pseudo-chromosomes contain one or both telomeric repeats (**Table S1c)**, and thus we conclude that all *Verticillium* strains have eight chromosomes, and that this number doubled in *V. longisporum*. Based on the inter-chromosomal Hi-C interaction signals, we identified a single region with high inter-chromosomal contacts for each of the pseudo-chromosomes (**Fig. S5; Table S1d**), indicating that these are the centromeres in the different *Verticillium* species. The average centromere size in *Verticillium* is ∼80 kb, yet we observed significant differences between the species (**Fig. 5b; Figs. S6a-b**). Centromeres within the Flavexudans clade are similarly sized and significantly smaller than the genus-wide average. By contrast, *V. dahliae* and *V. longisporum* centromeres are significantly larger.

**Figure 5.**
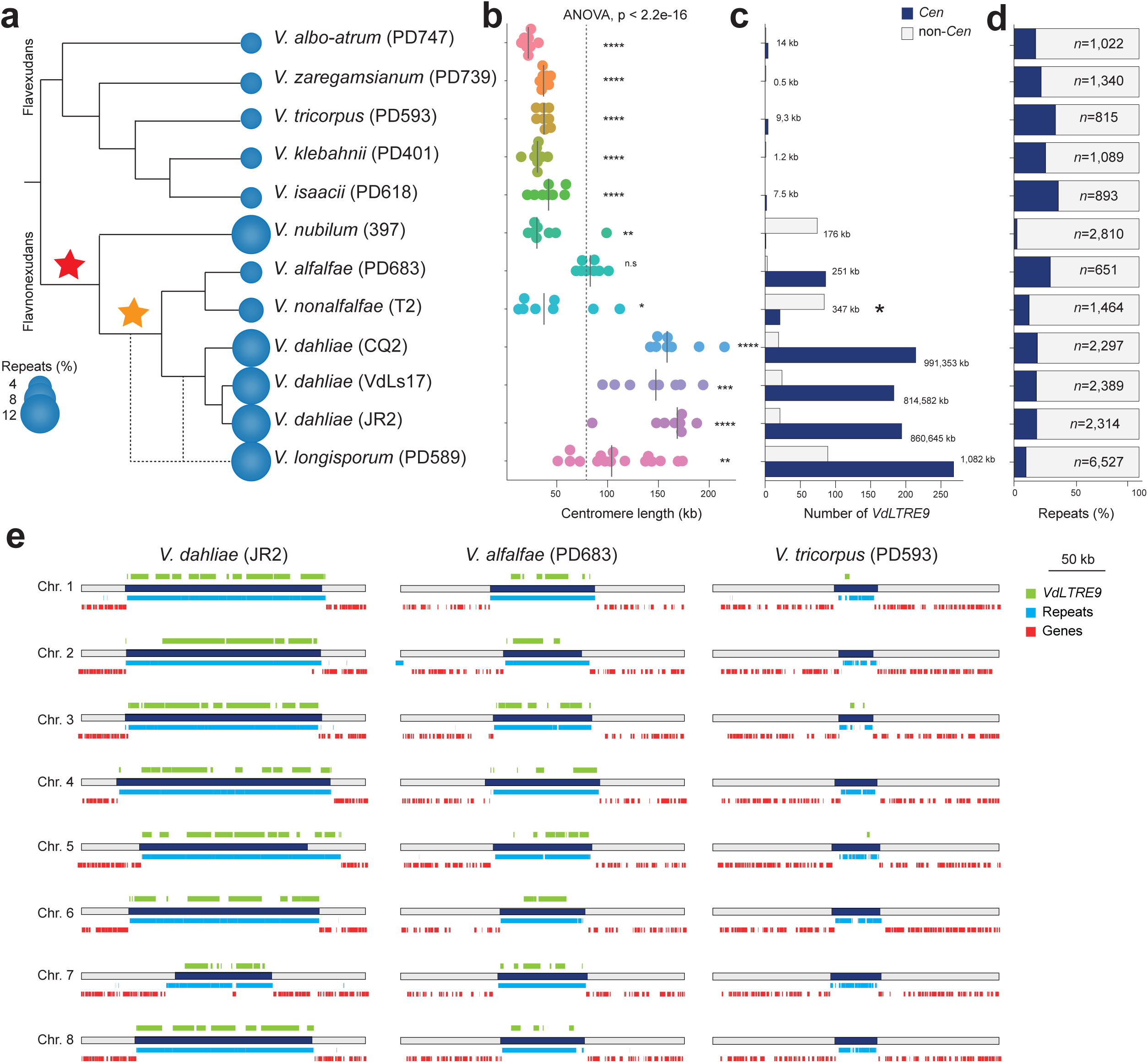
Evolution of centromeres in the genus *Verticillium.* (a) Relationship of the ten members of the genus *Verticillium*. The predicted repeat content for each of the genomes is indicated (see Table S3 for details). The red star indicates the acquisition of *VdLTRE9* in the Flavnonexudans clade while the yellow star indicates the recruitment of *VdLTRE9* into centromeres. (b) Comparison of estimated centromere lengths (in kb) in the different *Verticillium* spp. Each dot represents a single centromere and the line represents the median size. (c) The number of (partial) *VdLTRE9* matches identified in centromeres (*Cen*; dark blue) and across the genome (non-*Cen*; light grey). The asterisk indicates the high number of *VdLTRE9* elements in unassigned contigs for *Verticillium nonalfalfae* strain T2 (see text for details). (d) Proportion of predicted repeat content localized at centromeres (*Cen*; dark blue) and across the genome (non-*Cen*; light grey). (e) Schematic overview of the eight centromeric regions (250 kb) in *Verticillium dahliae* strain JR2, and *Verticillium alfalfae* strain PD683 and *Verticillium tricorpus* stain PD593 as representatives for clade Flavnonexudans and clade Flavexudans, respectively. The centromeres are indicated by dark blue bars. The predicted genes (red) and repeats (light blue) are shown below each centromere, and location of (partial) *VdLTRE9* matches (light green) are shown above each centromere. Global statistical differences for the centromere sizes was calculated using one-way ANOVA, and differences for each species compared to the overall mean were computed using unpaired T-tests; p-values < 0.0001: ****, p-values < 0.001: ***, p-values < 0.01: **, p-values < 0.05: *.

We subsequently assessed whether *VdLTRE9* defines centromeres in the other *Verticillium* species besides *V. dahliae* as well. Interestingly, *VdLTRE9* is abundant at centromeres in the allodiploid *V. longisporum* and in *V. alfalfae*, but fewer (21) or no *VdLTRE9* copies were identified at centromeres in *V. nonalfalfae* and *V. nubilum*, respectively (**Fig. 5c; Fig. S6c-d**). Similarly, only few or no (partial) matches of *VdLTRE9* consensus could be identified in the genomes of the Flavexudans species (**Fig. 5c; Fig. S6-7; Table S1e**). Collectively, these findings demonstrate that *VdLTRE9* is specific to Flavnonexudans species and has likely been recruited to the centromere only after the divergence of *V. nubilum* (**Fig. 5a; Fig. S6-7**).

Since *VdLTRE9* occurs only in few *Verticillium* species, we assessed to which extent other repetitive elements contribute to centromere organization. We analyzed the repeats identified by *de novo* repeat predictions for each of the *Verticillium* species. Centromeres in all species are AT- and repeat-rich (**Fig. S6a-b**), and some repeats occur in high frequency or nearly exclusively at centromeres in species that lack *VdLTRE9* (**Table S1e**). However, in contrast to *VdLTRE9*, these repeats cover only a minority (typically less than 10%) of the centromeres (**Table S1e**). Sequence similarity-based cluster analyses of the *de novo* repeat consensus sequences revealed that divergent repeat families contribute to *Verticillium* centromere organization (**Fig. S8**). Thus, in contrast to *VdLTRE9* in most Flavnonexudans species, we could not identify any additional repeat family as a hallmark of centromeres in other *Verticillium* species.

### Centromeres contribute to *Verticillium* karyotype evolution

We previously used fragmented genome assemblies to identify chromosomal rearrangements during *Verticillium* evolution (8, 35, 40). We hypothesize that centromeres might have contributed to these chromosomal rearrangements. To identify genome rearrangements and to trace centromeres during *Verticillium* evolution, we used the pseudo-chromosomes of the haploid *Verticillium* species to reconstruct ancestral chromosomal configurations using AnChro (**Fig 6a**) (56). We reconstructed all potential ancestors that predominantly had eight chromosomes and ∼8,000 genes (**Figs. S9a-b**), yet the number of ancestral chromosomes and genes varied when approaching the last common ancestor (**Figs. S9a-b**). By balancing the number of reconstructed chromosomes and genes, we identified a single most parsimonious ancestral genome with eight chromosomes and ∼8,500 genes (**Fig. 6a; Fig. S9c**), except for the last common ancestor within the clade Flavexudans clade that had eight major chromosomes and two additional ‘chromosomes’ with only six and two genes (**Fig. S9d**). As these two smaller ‘chromosomes’ likely do not represent genuine chromosomes, we conclude that all of the ancestral genomes, similar to the extant haploid *Verticillium* genomes, had eight chromosomes (**Fig. 6a**). Confirming our previous report (40), we observed in total 198 chromosomal rearrangements (124 inversions and 74 translocations) (**Fig. 6a**). The number of chromosomal rearrangements is lower than previously recorded and we did not observe any chromosomal fusion or fission events, which is likely the result of the drastically improved genome assemblies, but the rearrangement signal on each branch is sufficient to nevertheless recapitulate the known *Verticillium* species phylogeny (**Fig. S9e**). Importantly, we observed 17 genomic rearrangements that occurred at, or in close proximity (within ∼15 genes up or downstream) to, centromeres, both in extant *Verticillium* species as well as in the ancestors (**Fig. 6**). For example, at the branch from the last common ancestor (VA; **Fig. 6a**) to the ancestor of the clade Flavexudans (B1; **Fig. 6a**), two centromere-associated translocations (between the ancestral chromosome 2 and 6) led to the formation of two rearranged chromosomes. In total, we observed that five out of the eight ancestral centromeres were associated with a chromosomal rearrangement at one point during evolution (**Fig. 6a**). Nevertheless, comparisons of protein-coding genes that flank centromeres show that these are syntenic in most extant species. Similarly, none of the recent chromosomal rearrangements observed between *V. dahliae* strains is associated with centromeres (**Figs. 4a-b, 6a**). Thus, while chromosomal rearrangements involving centromeres occurred during evolution, they do not account for the majority of the karyotype variation between extant *Verticillium* species.

**Figure 6.**
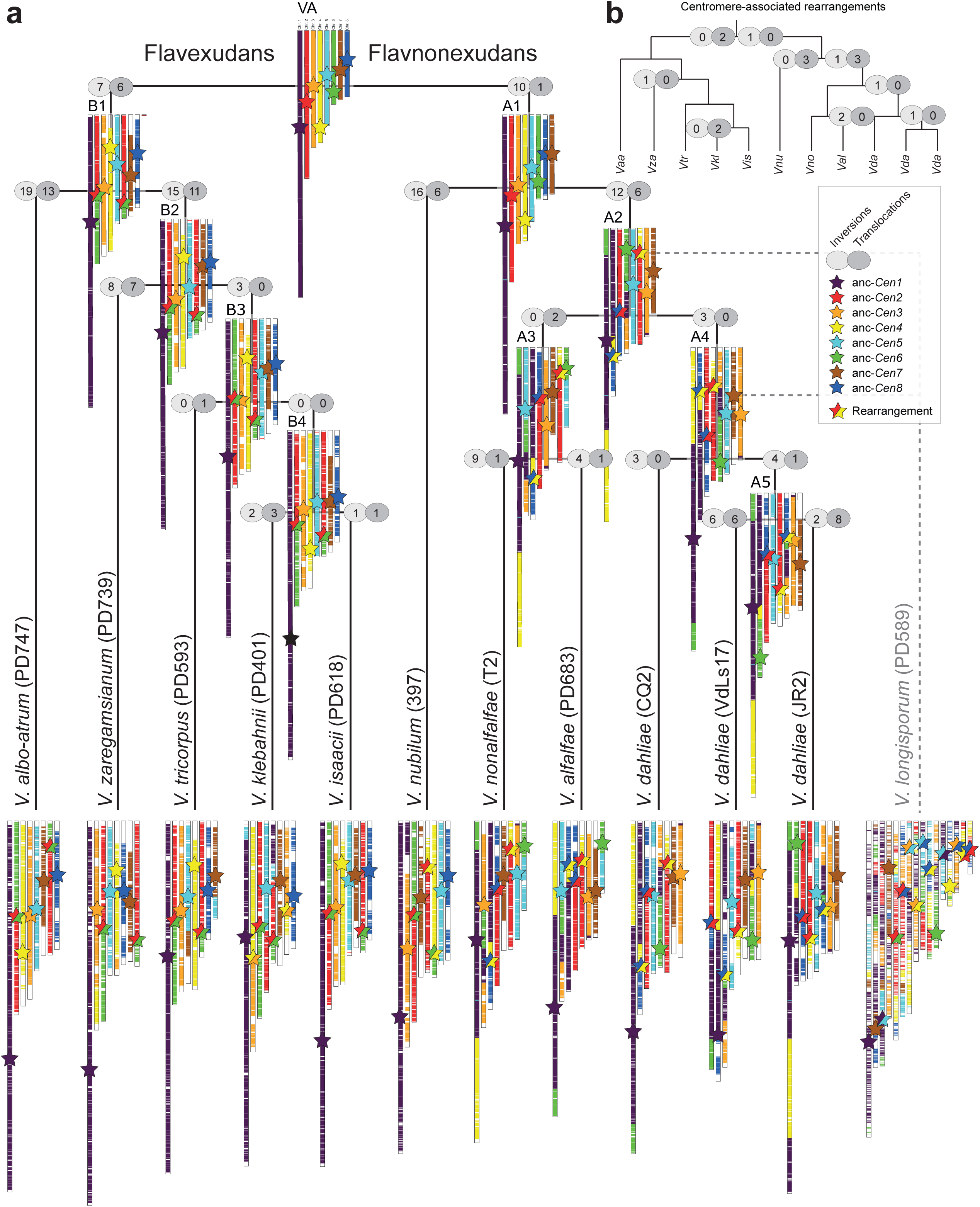
Centromeres contribute to karyotype evolution in *Verticillium.* (a) Relationship of the ten members of the genus *Verticillium*. The allodiploidization event forming *V. longisporum* is indicated by dashed lines (38, 57). The chromosomal evolution within the haploid members of the genus was reconstructed using AnChro (56). The chromosomal structure of the nine species is shown in relation to the last common ancestor of the genus. The approximate locations of the centromeres are indicated by stars. The number of chromosomal rearrangements (inversions and translocations) are displayed for each branch, and centromeres that co-localize in proximity to chromosomal rearrangements are highlighted by two-colored stars. (b) The number of major chromosomal rearrangements that occurred at, or in close proximity of, centromeres are shown along the branches depicting the *Verticillium* species phylogeny shown in (a).

## DISCUSSION

Centromeric regions are among the most rapidly evolving genomic regions (13–16, 29), yet centromere evolution has only been systematically studied in few fungi (11, 12, 16, 29). Here, we took advantage of the fungal genus *Verticillium* and used a combination of genetic and genomic strategies to identify and characterize centromere organization and evolution. *Verticillium* centromeres are characterized as large regional centromeres that are repeat-rich and embedded in heterochromatin. We furthermore show that centromeres contribute to the karyotype evolution of *Verticillium*. Finally, we demonstrate that *VdLTRE9* is a hallmark of centromeres in some *Verticillium* species, while species that lack *VdLTRE9* display a divergent repeat content.

Centromeres in fungi, plants, and animals co-localize within the nucleus (15, 51–55, 58), a phenomenon that can be exploited for their identification (51, 52). Here, we used Hi-C to first establish chromosome-level genome assemblies and subsequently identify centromeres in every *Verticillium* species, and we demonstrate that centromere locations are in agreement with CenH3-binding. While we obtained chromosome-level genome assemblies for all species, Hi-C scaffolded genome assemblies could still contain partially collapsed repeats and assembly gaps, in particular for short-read assemblies (59). With the exception of *V. nonalfalfae*, we observed only few sequencing gaps and no evidence that would point to collapsed repeats at centromeres, suggesting that the inferred centromeres are of high quality. *Verticillium* centromere sizes differ, which is likely not driven by assembly artefacts, and centromeres in most *Verticillium* species are larger than in *Z. tritici* (27), *C. neoformans, M. oryzae*, or *Fusarium graminearum* (13, 16, 29), yet smaller than in *N. crassa* (25). Species of the Flavexudans clade typically encode fewer repeats than species of the clade Flavnonexudans clade (32, 40, 60), and *V. nubilum*, *V. longisporum*, and *V. dahliae* are particularly rich in repeats when compared with other *Verticillium* species (32, 39–41, 60). Thus, increased centromere sizes positively correlate with overall increased repeat contents.

Using fragmented genome assemblies, we previously identified chromosomal rearrangements during *Verticillium* evolution (8, 35, 40) that were thought to have contributed to genetic diversity and adaptation in the absence of sexual recombination (7, 35, 40). Chromosome-level genome assemblies for an entire genus enabled unprecedented analyses of the karyotype evolution over longer evolutionary timescales. Here, we observed extensive chromosomal rearrangements and provide evidence that some rearrangements at centromeres contributed to karyotype evolution, most of which occurred early during the divergence of *Verticillium*. Chromosomal rearrangements at centromeres occur in the fungal yeasts *Candida*, *Cryptococcus,* and *Malassezia* (11, 12, 61), and synteny breakpoints have been identified between mammals and chicken (62), suggesting that centromeres often contribute to karyotype evolution. The emergence of chromosomal rearrangements at centromeres could be facilitated by their repeat-rich nature (11, 12). For example, centromeres in *Malassezia* are enriched with an AT-rich motif that could facilitate replication fork stalling, which leads to double strand DNA breaks (11). Repeats localized outside of centromeres in *V. dahliae* contribute to chromosomal rearrangements (8), and thus it seems plausible that centromeric repeats similarly contribute to chromosomal rearrangements. Chromosomal rearrangements often do not only lead to changes in chromosome organization but also in chromosome number (11, 12). While we observed chromosomal rearrangements, all extant and ancestral genomes contained eight chromosomes, suggesting that eight chromosomes are a stable configuration for all *Verticillium* species.

Centromere position and function are thought to be driven by the protein complement (e.g. CenH3 localization) and by heterochromatin formation rather than by specific DNA sequences (13, 15, 63). In *V. dahliae*, we observed the co-occurrence of CenH3 with H3K9me3 and DNA methylation. This suggests that DNA methylation, as previously reported in *N. crassa* and in *C. neoformans* (16, 25), is also a feature of centromeric DNA in *V. dahliae*. Co-localization of CenH3 with H3K9me2/3 and DNA methylation has been reported for *N. crassa* (25) and *C. neoformans* (16). In contrast, H3K9me3 and H3K27me3 are absent from centromeres in *Z. tritici* (27). H3K4me2 borders most centromeres in *Z. tritici* (27), and is associated with centromeres in *S. pombe* and some animals and plants (64–67). H3K4me2 has not been observed at centromeres in most fungi, including *V. dahliae*, and in the oomycete *P. sojae* (30). Changes in heterochromatin in *N. crassa* leads to altered CenH3 positioning (25), suggesting that heterochromatin is similarly required for centromere maintenance and function in *V. dahliae*. Elevated AT-levels in repeat-rich heterochromatic regions can be caused by RIP mutations (15, 25, 26, 42). RIP-like mutations have been previously reported in some repeats in *V. dahliae* (36, 44), and we observed strong RIP signals at centromeres. Due to its presumably asexual nature (7), the occurrence of RIP in *V. dahliae* is controversial (8, 43, 44). Noteworthy, mutational signatures resembling RIP have recently been observed in *Z. tritici* propagated through mitotic cell divisions, pointing to the existence of a mitotic version of a RIP-like process (42). Thus, we conclude that RIP was an active process in *V. dahliae* at some point in evolution, or that RIP-like processes outside of the sexual cycle occur in *V. dahliae*.

Centromeres are often enriched for a variety of different retrotransposons and other repetitive elements (15, 16, 25, 29, 30). We similarly observed that centromeres in all *Verticillium* species are repeat-rich. Repeats and their remnants identified at centromeres typically also occur outside of centromeres, as observed in *M. oryzae* (29) and *N. crassa* (25). Strikingly, we observed that a single repetitive element, *VdLTRE9*, is strongly associated with centromeres in some *Verticillium* species, which to our knowledge, has only been observed in the fungus *Cryptococcus* where centromeres contain six retrotransposons (*Tcn1*-*6*) that nearly exclusively occur at centromeres (16). Similarly, centromeres of the oomycete plant pathogen *Phytophthora sojae* contain multiple types of repeats, but they are enriched for a single element called CoLT (*<u>Co</u>pia*-<u>L</u>ike <u>T</u>ransposon) (30). The strong associations of specific repeats to centromeres could directly or indirectly link these elements to centromere function. Functional centromeres as observed here are also heterochromatic and contain CenH3. AT-rich repetitive elements can direct heterochromatin formation via DNA methylation and H3K9me3 deposition in *N. crassa* (45, 68), a phenomenon that can also occurs at repeats outside of centromeres (45). Heterochromatin occurs at centromeres but also at repeat-rich regions outside of centromeres in *V. dahliae*, thus the repeat-rich nature of centromeres is likely not sufficient to direct CenH3 deposition. In *S. pombe* heterochromatin formation is directed by short interfering RNAs (siRNA) derived from flanking repetitive elements via RNAi (69, 70), and RNAi and heterochromatin mediate CenH3 localization at centromeres (71, 72). RNAi is also important for centromere maintenance and evolution in *Cryptococcus*, as RNAi deficient species have smaller centromeres than RNAi proficient ones (16). Interestingly, centromere-specific elements (*Tcn1-6*) in RNAi proficient species are typically full-length elements while only remnants can be found in RNAi deficient species, which could be caused by recombination between elements (16). In *Verticillium*, centromere size differences correlate with increase of repeat content and the recruitment of *VdLTRE9*, which is highly fragmented and likely non-active. Furthermore, even though key components of the RNAi machinery exist in at least some *Verticillium* species (73), we know very little about its biological functions. Similarly, to *C. neoformans*, we observed no transcriptional activity of *VdLTRE9* or any other repeat at centromeres, but it is unclear if this silencing is mediated by RNAi, is a consequence of their heterochromatic nature, is due to their fragmentation, or a combination of these. Ultimately, unravelling how specific elements contribute to centromere identify necessitates future experiments. *VdLTRE9* occurs only in some *Verticillium* species and has likely been recruited to centromeres subsequent to the divergence of *V. nubilum*. Conversely, these observations raise further questions on the roles of repeats and mechanisms of centromeric identity in species without *VdLTRE9*. Repeats are important drivers of *Verticillium* genome evolution and function (8, 36), and here we highlight their contributions to centromere diversity within the fungal genus *Verticillium*. Our analyses provide the framework for future research into the diversity or convergence of mechanisms establishing centromere identity and functioning in fungi.

## MATERIAL & METHODS

### Construction of *Verticillium dahliae* transformants expressing FLAG-tagged CenH3

CenH3 and H3 homologs were identified in the predicted proteomes of *V. dahliae* strain JR2 (32) and selected other fungi through a BLAST sequence similarity search (blastp v2.9.0+; default settings, e-value cutoff 1e-20) (74, 75) using the *N. crassa* CenH3 (Q7RXR3) and H3 (P07041) sequences as queries. Missing homologs of CenH3 or H3 were identified using manual BLAST (tblastn v2.9.0+; default settings) (74, 75) and exonerate (v2.2.0; default settings) (76) searches against the genome sequences. Protein sequences of selected CenH3 and H3 proteins were aligned using mafft (v7.271; default settings, LINSi) (77). A phylogenetic tree was inferred with maximum-likelihood methods implemented in IQ-tree (v1.6.11) (78) and robustness was assessed by 1,000 rapid bootstrap replicates.

To construct the N-terminally FLAG-tagged CenH3 strain of *V. dahliae,* a recombinant DNA fragment was constructed into the binary vector PRF-HU2 (79) or PRF-GU2 for homologous recombination. The CenH3 locus, from *V. dahliae* strain JR2, was amplified as 3 fragments with overlapping sequences (**Table S1f**). The 5’ most fragment containing the promoter was amplified using primers A + B, the ORF with primers C+D, the Hyg promoter and ORF with primers E+F, and the 3’ end of the CenH3 locus with primers G+H. The four fragments were combined by overlap PCR using primers A + H and cloned into a *Psp*OMI and *Sp*hI linearized vector using Gibson Assembly. The vector construction was confirmed by Sanger sequencing. Vectors were transformed to *Verticillium* with *Agrobacterium-*mediated transformation (80). Correct homologous recombination and replacement at the *CenH3* locus was verified by PCR amplification using primer I+J (**Fig. S1b**, **Table S1f**). Correct translation of the recombinant protein was assessed using Western analyses with anti-FLAG antibody (**Fig. S1c**). Briefly, proteins were extracted from 5-day old cultures grown in 100 ml Potato Dextrose Broth at 22°C with continuous shaking at 120 rpm. Mycelium was collected by straining over a double layer of miracloth and subsequently snap-frozen in liquid nitrogen and ground with a mortar and pestle using liquid nitrogen. Approximately 0.3 g of ground mycelium was resuspended in 600 µL protein extraction buffer (50 mM HEPES pH 7.5, 150 mM NaCl, 1 mM EDTA, 1% glycerol, 0.02% NP-40, 2 mM Phenylmethanesulfonyl fluoride (PMSF), 100 µM Leupeptin, 1 µg/mL Pepstatin), briefly vortexed, incubated on ice for 15 min and centrifuged at 4°C at 8,000 g for 3 min. The supernatant was collected by transferring 20 µL to a new tube to serve as the input control and the remaining ∼500 µL was transferred to a fresh microcentrifuge tube with 15 µL of Anti-FLAG M2 affinity gel (catalog number A2220, Sigma-Aldrich, St. Louis, Missouri, United States) and incubated while rotating at 4°C for 1 h. Samples were centrifuged at 5,000 g, 4°C for 3 min, after which the supernatant was discarded. Samples were washed with 500 µL of lysis buffer, and the centrifugation and washing were repeated three times. Protein was eluted from the resin by adding 15 µL of lysis buffer, 20 µL of 2x Laemmli loading buffer (4% SDS, 20% glycerol, 0.004% bromophenol blue, 125 mM Tris HCL pH 6.8) and boiled at 95°C for 3 min. Protein samples were separated on a 12% polyacrylamide gel, and subsequently transferred to PVDF membranes, blocked in 5% BSA, washed twice in TBST, and incubated with 1:3500 anti-FLAG antibody (monoclonal anti-FLAG M2; Merck KGaA, Darmstadt, Germany).

### Chromatin immunoprecipitation followed by high-throughput sequencing (ChIP-seq)

For each *V. dahliae* genotype, one million spores were added to 100 ml Potato Dextrose Broth and incubated for 7 days at 22°C with continuous shaking at 120 rpm. Mycelium was collected by straining over a double layer of miracloth and subsequently snap-frozen in liquid nitrogen and ground with a mortar and pestle using liquid nitrogen. All ground material (0.5-1 gram per sample) was resuspended in 4 mL ChIP Lysis buffer (50 mM HEPES-KOH pH7.5, 140 mM NaCl, 1 mM EDTA, 1% Triton X-100, 0.1% NaDOC) and dounced 40 times in a 10 cm^3^ glass tube with tightly fitting pestle on 800 power with a RZR50 homogenizer (Heidolph, Schwabach, Germany), followed by five rounds of 20 seconds sonication on ice with 40 seconds of resting in between rounds with a Soniprep 150 (MSE, London, UK). Samples were redistributed to 2 mL tubes and pelleted for 2 min at maximum speed in a tabletop centrifuge. Supernatants were pooled per sample in a 15 mL tube together with 25 μL α-FLAG M2 magnetic beads (Sigma-Aldrich, St. Louis, Missouri, United States), incubated overnight at 4°C and continuous rotation. Beads were captured on a magnetic stand and washed with wash buffer (50 mM Tris HCl pH 8, 1 mM EDTA, 1% Triton X-100, 100 mM NaCL), high-salt wash buffer (50 mM Tris HCl pH 8, 1 mM EDTA, 1% Triton X-100, 350 mM NaCL), LiCl wash buffer (10 mM Tris HCl pH8, 1 mM EDTA, 0.5% Triton X-100, 250 mM LiCl), and TE buffer (10 mM Tris HCl pH 8, 1mM EDTA). Chromatin was eluted twice from beads by addition of 100 μL pre-heated TES buffer (100 mM Tris HCl pH 8, 1% SDS, 10 mM EDTA, 50 mM NaCl) and 10 minutes incubation at 65°C. 10 mg/mL 2 μL Proteinase K was added and incubated at 65°C for 5 hours, followed chloroform extraction. DNA was precipitated by addition of 2 volumes 100% ethanol, 1/10^th^ volume 3 M NaOAc pH 5.2 and 1/200^th^ volume 20mg/mL glycogen, and overnight incubation at −20°C.

Sequencing libraries were prepared using the TruSeq ChIP Library Preparation Kit (Illumina, city, country) according to the manufacterer’s instructions, but without gel purification and with use of the Velocity DNA Polymerase (BioLine, Luckenwalde, Germany) for 12 cycles of amplification for the FLAG-CenH3. H3K4me2 ChIP was performed as described previously (36), using an α-H3K4me2 antibody (#39913, ActiveMotif; Carlsbad, California, United States). Single-end (125 bp) sequencing was performed on the Illumina HiSeq2500 platform at KeyGene N.V. (Wageningen, the Netherlands).

### Chromatin confirmation capturing followed by high-throughput sequencing (Hi-C)

We determined the inter- and intra-chromosomal contact frequencies using Hi-C in *V. dahliae* strains CQ2, JR2, and VdLs17, as well as in *V. albo-atrum* strain PD747, *V. alfalfae* strain PD683, *V. isaacii* strain PD618, *V. klebahnii* strain PD401, *V. longisporum* strain PD589, *V. nonalfalfae* strain T2, *V. nubilum* strain 397, *V. tricorpus* strain PD593, and *V. zaregamsianum* strain PD739. For each strain, one million spores were added to 400 mL Potato Dextrose Broth and incubated for 6 days at 22°C with continuous shaking at 120 rpm. Mycelium was collected by straining over double layer miracloth and 300 mg (fresh weight) was used as input for generating Hi-C sequencing libraries with the Proximo Hi-C kit (Microbe) (Phase Genomics, Seattle, WA, USA), according to manufacturer’s instructions. Hi-C sequencing libraries of V*. dahliae* strains CQ2, JR2. and VdLs17 were paired end (2×125 bp) sequenced on the Illumina HiSeq2500 platform at KeyGene N.V. (Wageningen, the Netherlands). Hi-C sequencing libraries of the other *Verticillium* species were paired-end (2×150 bp) sequenced on the NextSeq500 platform at USEQ (Utrecht, the Netherlands).

### *In vitro* transcriptome profiling using RNA-seq

RNA sequencing of *V. albo-atrum* strain PD747, *V. isaacii* strain PD618, *V. klebahnii* strain PD401, *V. longisporum* strain PD589, *V. nonalfalfae* strain T2, *V. nubilum* strain 397, *V. tricorpus* strain PD593, and *V. zaregamsianum* strain PD739 as described previously (36). Single-end (50 bp) sequencing was performed on the BGISeq500 platform at BGI (BGI Hong Kong).

### Analyses of high-throughput sequencing data

High-throughput sequencing libraries (**Table S1a**) have been analyzed as follows: Illumina reads were quality-filtered and trimmed using trimmomatic (version 0.36) (81). Sequencing reads were trimmed and filtered by removing Illumina TruSeq sequencing adapters (settings seed mismatches 2, palindrome clip threshold 30, and simple clip threshold 10), removal of low-quality leading or trailing bases below quality 5 and 10, respectively, and 4-base sliding window trimming and cutting when average quality per base dropped below 15. Additionally, filtered and trimmed reads < 90 nt were removed from further analyses. Filtered and trimmed reads were mapped to the corresponding genome assembly with Bowtie2 (default settings) (82), and mapping files were converted to bam-format using samtools (v 1.8) (83). Genomic coverage was determined using deepTools (v3.4.1; bamCoverage) (84) by extending sequencing reads to 147 bp followed by RPGC normalization with a bin-size of 1,000 bp and smoothening of 3,000 bp. To assess between sample variability, we used deepTools (v3.4.1, plotPCA) (84) to generate principle component analyses. Furthermore, we employed deepTools (v3.4.1, multiBigwigSummary) (84) to summarize genomic coverages of over genes, repetitive elements, and genomic windows (5 kb windows with 500 bp slide). Genomic regions enriched for FLAG-CenH3 were identified using MACS2 (v2.1.1) (broad peak option; broad cutoff 0.0025) (85).

To determine DNA (cytosine) methylation, we utilized sequencing data of bisulfite treated genomic DNA previously generated for *V. dahliae* strain JR2 (36). Sequencing reads were mapped to the *V. dahliae* strain JR2 genome assembly as previously described (36). Subsequently, the number of reads supporting cytosine methylation in CG-context were extracted, and weighted CG-methylation levels were calculated over genes, repetitive elements, and genomic windows (5 kb window size with 500 bp slide) (86); weighted CG-methylation was defined as the sum of reads supporting cytosine methylations divided by the sum of all reads occurring at all CG sites in the respective regions. Sites with less than four reads were not considered.

To improve the genome assemblies of the *Verticillium* species, we mapped Hi-C sequencing reads to genome assemblies of *V. dahliae* strain CQ2, *V. albo-atrum* strain PD747, *V. alfalfae* strain PD683, *V. isaacii* strain PD618, *V. klebahnii* strain PD401, *V. longisporum* strain PD589, *V. nonalfalfae* strain T2, *V. nubilum* strain 397, *V. tricorpus* strain PD593, and *V. zaregamsianum* strain PD739 using Juicer (v1.6) with early stage setting (87). The contact matrices generated by juicer were used by the 3D de novo assembly (3D-DNA) pipeline (88) (v180922) with a contig size threshold of 1000bp to eliminate mis-joints in the previous assemblies and to generate improved assemblies. The genome assemblies were manually improved using Juicebox Assembly Tools (JBAT) (v1.11.08) (89) and improved genome assemblies were generated using the 3D-DNA post-review asm pipeline (88). Centromere locations were determined using a 1 kb-resolution contact matrix in JBAT, by identifying a region per chromosome that displays strong inter-chromosomal interactions, yet weak intra-chromosomal interactions (see Figure S12, S13).

To assess potential repeat collapses during genome assemblies at centromeric regions, we mapped previously generated short-read data *V. dahliae* strain JR2 and VdLs17, *V. albo-atrum* strain PD747, *V. alfalfae* strain PD683, *V. isaacii* strain PD618, *V. klebahnii* strain PD401, *V. longisporum* strain PD589, *Verticillium nonalfalfae* strain T2, *V. tricorpus* strain PD593, and *V. zaregamsianum* strain PD739 (39, 40, 90, 91) to the genome assemblies using BWA (v0.7.17; mem) (83). We first used bedtools (v2.29.2) (92) to identify few genomic regions with > 500x coverage. We then applied deepTools (v3.4.1, computeGCBias) (84) to compute GC biases of read depth across the genome, excluding the identified high coverage regions, and used deepTools (v3.4.1, correctGCBias) (84) to correct GC biases, which addresses known biases in sequencing library preparation to ensure even read coverage throughout the genome irrespective of their base composition (93). We used deepTools (v3.4.1, bamCoverage, bins 50 bp, CPM normalization) (84) to obtain the read coverage throughout the genome, excluding regions containing sequence assembly gaps (Ns). Assuming that collapsed repeats would lead to a local increase in read depth, we used the ratio of the average read coverage at the centromeres and outside of the centromere at each chromosome to correct the inferred centromere sizes. To further validate the genome assembly of regions identified as centromeres of *V. dahliae* strain JR2, the genome assembly was compared to the previously generated optical map (35) using MapSolver (v 3.2; OpGen, Gaithersburg, MD).

The transcriptional activity for genes and repetitive elements in *V. dahliae* strain JR2 was assessed *in vitro* (in Potato Dextrose Broth) using previously generated deep transcriptome datasets (36). To this end, single-end sequencing reads of three biological replicates were mapped to the *V. dahliae* strain JR2 genome assembly (32) using STAR (v2.4.2a; max. intron size 1 kb and outFilterMismatchNmax to 5) (94). The resulting mapped reads were summarized per genomic feature (gene or repeat) using summarizeOverlaps (95), converted to counts per million (cpm) mapped reads, and averaged over the three biological replicates.

### Sequence analyses of *Verticillium* genome assemblies, centromeres, repeat and gene content

Repetitive elements in the genomes of *V. dahliae* strains JR2, VdLs17 and CQ2 (32, 33) were identified as previously described (36). Briefly, repetitive elements were identified in each genome independently using a combination of LTRharvest (96) and LTRdigest (97) followed by identification of RepeatModeler. Identified repeats in the different *V. dahliae* strains were clustered into a non-redundant library that contained consensus sequences for each repeat family. The repeat library was manually curated and annotated using PASTEC (98) or by sequence similarity to previously identified and characterized repeat families (32, 44). Genome-wide occurrences of repeat families were determined using RepeatMasker (v 4.0.9; sensitive option and cutoff 250), and the output was postprocessed using ‘One code to find then all’ (99). We only considered matches to the repeat consensus library, and thereby excluded simple repeats and low-complexity regions.

*De novo* gene and repeat annotation for the Hi-C-improved *Verticillium* genome assemblies, and for *V. dahliae* strains JR2 and VdLs17 as a comparison was performed using the funannotate pipeline (100). Briefly, repetitive elements were first *de novo* identified using RepeatModeler and masked for gene prediction using RepeatMasker. Subsequently, gene prediction parameters were estimated using *in vitro* RNA-seq data (see above for details; exception: *V. alfalfae* for which no RNA-seq data was available, *V. nonalfalfae* for which publicly available RNA-seq data was used (90), and *V. dahliae* strain JR2 for which in addition to the *in vitro* RNA-seq data generated in this study, also previously generated *in vitro* (xylem sap and half-MS; (36)) as well as long-read nanopore cDNA data (101) was used). Based on the gene prediction parameters, gene prediction was performed with funannotate using a combination of *ab initio* gene predictors, consensus predictions were obtained using Evidencemodeler (v1.1.1) (102), and gene predictions were adjusted using information from the RNA-seq data. Repeat annotation for each genome assembly was based on the *de novo* repeat family consensus sequences obtained with funannotate. Genome-wide occurrences of these repeat families as well as previously defined repeat families for *V. dahliae* (see above) were determined using RepeatMasker (v 4.0.9; sensitive option and cutoff 250), and the output was postprocessed using ‘One code to find then all’ (99). *De novo* repeat families overlapping with centromeres in the different species were clustered using BLASTClust (v2.2.26; parameter ‘-S 60 -L 0.55 -b F -p F’), and subsequently visualized using Cytoscape (v.3.8.0) (103). Next to RepeatMasker, genome-wide occurrences of the previously determined *VdLTRE9* (32, 36) were identified by BLAST searches (blastn v2.9.0+; e-value cutoff 1e-5, no soft-masking and dust, fixed database size 10e6) (74, 75), and similarity between VdLTRE9 consensus sequences and the *de novo* predicted repeat families was established using BLAST (blastn, e-value cutoff 1e-5, query coverage > 50%, no soft-masking and dust, fixed database size 10e6).

Repeat and gene density (*V. dahliae* strain JR2 and VdLs17 based on previous gene annotation (101)), GC-content, and composite RIP index were calculated along the genome sequence using sliding windows (5 kb window with 500 bp slide). The composite RIP index (CRI) was calculated according to Lewis et al. (45). CRI was determined by subtracting the RIP substrate from the RIP product index, which are defined by dinucleotide frequencies as follows: RIP product index = TpA / ApT and the RIP substrate index = (CpA + TpG)/(ApC + GpT). Overlaps between different genomic features (for example repetitive elements over centromeric regions) was assessed using bedtools (v2.29.2) (92). Genome-wide data was visualized using R (104) with the packages ggplot2 (105), karyplotR (106), or Gviz (107), as well as EasyFig (108).

Whole-genome alignments between *V. dahliae* strains JR2, VdLs17, and CQ2 were performed using NUCmer, which is part of the MUMmer package (v 3.1; --maxmatch) (109). To remove short matches, we only considered alignments longer than 10 kb. Ancestral genome configurations were reconstructed using AnChro (56). We first determined the synteny relationships between all possible pairs of haploid *Verticillium* genomes and two outgroup genomes (*Plectosphaerella cucumerina* and *Sodiomyces alkalinus*) using SynChro with synteny block stringency (delta parameter) ranging from 2-5 (110). We then obtained all ancestors by calculating all possible pairs of genomes (G1 and G2) and outgroups (G3,..,G_n_) and by varying the delta’ (G1 and G2 comparisons) and delta” (G1/G3..G1/G_n_ and G2/G3..G2/G_n_ comparisons) parameters for AnChro. We additionally reconstructed all ancestors starting from the extent genomes in a sequential approach with multiple successive cycles of SynChro and AnChro (delta parameters varied between 2-5). For each ancestor, we chose the optimal reconstructed by the delta parameter combination (delta’ and delta”) that minimizes the number of reconstructed chromosomes and rearrangements and at the same time maximizes the number of genes, both guided by the most commonly observed number of chromosomes and genes in all rearrangements. We obtained the number of large-scale rearrangements between reconstructed ancestral genomes and the extent *Verticillium* genomes using ReChro with a delta parameter of 1 (56). The relationship between chromosomes of the reconstructed ancestors and the extent species in relationship to the common ancestor is generated with SynChro with a delta parameter of 1 (110). A species phylogeny that uses synteny relationships computed by SynChro (see above) as informative character between the *Verticillium* genomes and the outgroup genomes was reconstructed using PhyChro (111).

### Data availability

ChIP-seq and Hi-C data were submitted to the Short Read Archive (SRA) under the accession PRJNA641329 (**Table S1a**).

## ACKNOWLEDGMENTS

Work in the laboratories of M.F.S and B.P.H.J.T. is supported by the Research Council Earth and Life Sciences (ALW) of the Netherlands Organization of Scientific Research (NWO). Furthermore, B.P.H.J.T. would like to acknowledge the Deutsche Forschungsgemeinschaft (DFG, German Research Foundation) under Germanýs Excellence Strategy – EXC 2048/1 – Project ID: 390686111. This work was supported in part by a European Molecular Biology Organization postdoctoral fellowship (EMBO, ALTF 969-2013) and Human Frontier Science Program Postdoctoral Fellowship (HFSP, LT000627/2014-L) to D.E.C. A portion of the work was also carried out in the laboratory of D.E.C. under USDA-NIFA-PBI grant 2018-67013-28492. We thank Utrecht Sequencing Facility for providing sequencing service and data. Utrecht Sequencing Facility is subsidized by the University Medical Center Utrecht, Hubrecht Institute, Utrecht University, and The Netherlands X-omics Initiative (NWO project 184.034.019).

## SUPPLEMENTARY MATERIAL

**Figure S1** – (**a**) Phylogenetic analyses of the canonical H3 and the centromeric-specific CenH3 in *Verticillium dahliae* (strain JR2) and other fungal genomes. (**b-c**) Transformation of the coding sequence of N-terminally FLAG-tagged CenH3 directed by its native promoter at the *CenH3* locus in *Verticillium dahliae* strain JR2. (**b**) Correct homologous recombination and replacement at the *CenH3* locus was verified by PCR amplification was assessed using PCR and (**c**) Correct translation of the recombinant protein was assessed using Western Blot analyses with anti-FLAG antibody. (**d**) Sequencing read coverage (RPGC normalization in 1 kb bins with 3 kb smoothening) from ChIP-seq experiments using FLAG-tag antibodies on two independent transformants of *Verticillium dahliae* strain JR2 that express FLAG-tagged CenH3 and the wild-type strain are mapped to the eight chromosomes of *V. dahliae* strain JR2 (32). Gene (red) and repeat (blue) density are shown below each chromosome. (**e**) Principal component analysis of the four FLAG-tag ChIP-seq samples (two wild-type and two CenH3-FLGA). (**f**) Comparison of the centromeric regions with the identified centromeres highlighted as blue block in the genome assembly of *Verticillium dahliae* strain JR2 with a previously generated optical map (35). Vertical lines display corresponding (*in silico*) restriction sites and their alignment.

**Figure S2** – Schematic overview of the eight chromosomes of *Verticillium dahliae* strain JR2 displaying different heterochromatin-associated chromatin modifications (mC, H3K9me3, and H3K27me3) in relation to the centromeres. The different lanes display the CenH3-FLAG ChIP-seq read coverage (RPGC normalization in 1 kb bins with 3 kb smoothening), the repeat-density, the GC-content, the CRI as well as the weighted cytosine methylation (all summarized in 5 kb windows with 500 bp slide), and the normalized H3K9me3 and H3K27me3 ChIP-seq read coverage (RPGC normalization in 1 kb bins with 3 kb smoothening).

**Figure S3** – (**a**) Boxplot displaying the composite RIP index (CRI) of C to T in CA recorded in genomic windows (5 kb, 500 bp slide), per gene, per annotated repeat, and per window overlapping with the CenH3-enriched centromeres. Statistical differences for the indicated comparisons were calculated using the one-sided non-parametric Mann-Whitney test; p-values < 0.001: ***. (**b**) Summary of H3K4me2 (green), H3K9me3 (red), and H3K27me3 (orange) normalized ChIP-seq read coverage (RPGC normalization in 1 kb bins and 3 kb smoothening) in genomic bins (2.5%) across the chromosomal arms of the eight chromosomes of *Verticillium dahliae* strain JR2 (divided into 2.5% bins) and the centromeric regions (divided into 10% bins). The dots indicate the average ChIP-seq coverage and the whiskers indicate ± 1.5 times the interquartile range. (**c-e**) Boxplots displaying the (**c**) weighted methylation levels (CG context), (**d**) the composite RIP index, and (**e**) the expression in PDB growth medium (counts per million) for repetitive elements belonging to ten repeat families identified in the eight centromeres in *Verticillium dahliae* JR2.

**Figure S4** (**a-c**) Whole-genome alignments between the eight chromosomes of (**a**) *Verticillium dahliae* strains JR2 and VdLs17 (32), (**b**) *V. dahliae* strains CQ2 and JR2 (32, 33), and (**c**) *V. dahliae* strains CQ2 and VdLs17 (32, 33). (**d-e**) Schematic overview of the genome assemblies of *Verticillium dahliae* strains (**d**) VdLs17 and (**e**) CQ2. The individual lanes show the GC content, the gene (red) and repeat (blue) density (all summarized in 5 kb windows with 500 bp slide), and the location of the centromere associated *VdLTRE9*. (**f**) Synteny analyses of the eight chromosomes of *V. dahliae* strains JR2 and CQ2. Schematic overview of the eight chromosomes of *V. dahliae* strain JR2 (left) and the corresponding syntenic regions in *V. dahliae* strains CQ2 (right). Centromeres are indicated by stars, and syntenic centromeres of *V. dahliae* strain CQ2 are colored according to *Cen1*-*8* of *V. dahliae* strain JR2.

**Figure S5** – Hi-C contact matrix showing the interaction frequencies between genomic regions in (**a**) *V. nonalfalfae* (T2), (**b**) *V. alfalfae* (PD683), (**c**) the allodiploid *V. longisporum* (PD589), (**d**). *V. nubilum* (397), (**e**) *V. albo-atrum* (PD747), (**f**) *V. zaregamsianum* (PD739), (**g**) *V. tricorpus* (PD593), (**h**) *V. klebhanii* (PD401), and (**i**) *V. isaacii* (PD618). Regions of high inter-chromosomal interaction frequencies are indicative of centromeres and are highlighted by arrow heads, and the blue line indicated boundaries between the pseudo-chromosomes.

**Figure S6** – (**a-b**) Comparison of normalized read coverage and corrected centromere lengths for *Verticillium* species for which short-read data is available. (**a**) Counts per million mapped reads (CPM) normalized read coverage was calculated for GC-biased corrected short-read libraries in 50 bp genomic windows, excluding regions containing assembly gaps (Ns). Genomic windows are summarized in boxplots (outliers not shown) by genomic location, centromeric regions (*Cen*, blue) and non-centromeric regions (non-*Cen*, grey). (**b**) Centromeric lengths inferred by Hi-C data were ‘corrected’ based on the ratio of normalized read depth between centromeres and non-centromeric regions per chromosomes. Differences for each species compared to the overall mean were computed using unpaired T-tests; p-values < 0.0001: ****, p-values < 0.001: ***, p-values < 0.01: **, p-values < 0.05: *. (**c**) The number of BLASTn matches of the *VdLTRE9* consensus element to the genomes of the *Verticillium* species separated by their genomic location, centromeric regions (*Cen*, blue) and non-centromeric regions (non-*Cen*, grey). The overall number of base pairs (bp) covered by the BLASTn matches in each genome sequence is indicated. The asterisk denotes the high number of *VdLTRE9* matches to unassigned, non-*Cen* regions in the genome assembly of *Verticillium nonalfalfae* (T2). (**d**) The number of repetitive element matches identified by RepeatMasker for each *Verticillium* species based on species/strain-specific repeat libraries generated by RepeatModeler separated by their genomic location, centromeric regions (*Cen*, blue) and non-centromeric regions (non-*Cen*, grey). (**e**) GC-content of the *Verticillium* genomes in 50 bp windows and separated by their genomic location, centromeric regions (*Cen*, blue) and non-centromeric regions (non-*Cen*, grey). (∫) The repeat content of centromeric regions in percent covered sequences in the different *Verticillium* species. Each data point summarized in the boxplot is the repeat content per centromere.

**Figure S7** – Schematic overview of the centromeric regions (250 kb) in (**a**) *Verticillium dahliae* strain JR2, in (**b**) species belonging to clade Flavnonexudans, and in (**c**) species belonging clade Flavexudans. The centromeres are indicated by dark grey bars. The predicted genes (black) and repeats (blue) are shown below each centromere, and location of *VdLTRE9* (partial) matches (light green) are shown above each centromere. Repeats that share sequence similarity (BLASTn) to the *VdLTRE9* consensus sequence are shown above each centromere (dark green).

**Figure S8** – Sequence comparisons of *de novo* repeat families identified with RepeatModeler and RepeatMasker in the genome assemblies of the different *Verticillium* species. Individual repeat family consensus sequences were clustered using BLASTClust. (**a**) Relationships between different repeat family consensus sequences are displayed as connected graphs. The sub-graph with the consensus sequences with similarity to *VdLTRE9* is highlighted in yellow. (**b**) The presence/absence matrix indicates the occurrences of different repeat families in the analyzed *Verticillium* species (black present, white absent). The cluster containing consensus sequences with similarity to *VdLTRE9* is highlighted.

**Figure S9** – Reconstruction of ancestral genomes within the genus *Verticillium* with AnChro (56). The number of (**a**) chromosomes and (**b**) genes predicted by all potential ancestral reconstructions using different combinations of genomes and stringency parameters. The phylogenetic tree in (**a**) depicts the relationships between *Verticillium* species and the abbreviations used for the ancestors. The inlays display boxplots to summarize the number of (**a**) chromosomes and (**b**) genes per ancestral reconstruction. (**c**) The number of chromosomes and genes of the chosen ‘optimal’ reconstruction for each of the internal ancestors. (**d**) The number of genes per chromosome for each of the reconstructed ancestor and the extant *Verticillium* species. The star highlights the reconstruction for the B1 ancestor that had ten chromosomes, but with two chromosomes with six and two genes. (**e**) Reconstruction of the *Verticillium* species phylogeny based on synteny relationship using PhyChro (111).

**Table S1 –** (a) Overview of the different *Verticillium* sequencing libraries used in this study. (b) Position of the individual centromeric regions inferred by Hi-C inter-chromosomal interaction frequencies and the overlap (in kb) with CenH3-enriched regions and the centromere associated *VdLTRE9* in *Verticillium dahliae* JR2, VdLs17, and CQ2. (c) Overview of the different *Verticillium* genomes assembled using Hi-C interactions. (d) Position, length, and number of assembly gaps (Ns) of the individual centromeric regions inferred by Hi-C inter-chromosomal interaction in V*erticillium nonalfalfae* (T2), *Verticillium alfalfae* (PD683), the allodiploid *Verticillium longisporum* (PD589), *Verticillium nubilum* (397),*Verticillium albo-atrum* (PD747), *Verticillium zaregamsianum* (PD739), *Verticillium tricorpus* (PD593), *Verticillium klebhanii* (PD401), and *Verticillium isaacii* (PD618). (e) The number of *de novo* repeat consensus sequences identified within and outside of centromeric regions in the *Verticillium* species. Only consensus elements with > 5 matches in centromeric regions are displayed. Note that the consensus names between species/strains are not comparable. (f) The primers used for cloning the CenH3 FLAG tag in *Verticillium dahliae* strain JR2

